# P-TEFb is degraded by Siah1/2 in quiescent cells

**DOI:** 10.1101/2021.12.30.474394

**Authors:** Fang Huang, Yongmei Feng, B. Matija Peterlin, Koh Fujinaga

**Affiliations:** Departments of Medicine, Microbiology and Immunology, University of California, San Francisco, San Francisco, CA 94143, USA; Cancer Center, Sanford Burnham Prebys Medical Discovery Institute, La Jolla, CA 92037, USA

## Abstract

P-TEFb, composed of CycT1 and CDK9, regulates the elongation of transcription by RNA polymerase II. In proliferating cells, it is regulated by 7SK snRNA in the 7SK snRNP complex. In resting cells, P-TEFb is absent, because CycT1 is dephosphorylated, released from CDK9 and rapidly degraded. In this study, we identified the mechanism of this degradation. We mapped the ubiquitination and degradation of free CycT1 to its N-terminal region from positions 1 to 280. This region is ubiquitinated at six lysines, where E3 ligases Siah1 and Siah2 bind and degrade these sequences. Importantly, the inhibition of Siah1/2 rescued the expression of free CycT1 in proliferating as well as resting primary cells. We conclude that Siah1/2 are the E3 ligases that bind and degrade the dissociated CycT1 in resting, terminally differentiated, anergic and/or exhausted cells.

## INTRODUCTION

Eukaryotic transcription by RNA polymerase II (RNAPII) consists of initiation, promoter clearance, elongation and termination. After alternative cleavage and polyadenylation, the single strand RNA is transported to the cytoplasm for translation (1–4). Phosphorylation of RNAP II’s C-terminal domain (CTD) at position 5 (Ser5) in tandemly repeated heptapeptides (Tyr-Ser-Pro-Thr-Ser-Pro-Ser) by cyclin-dependent kinase 7 (CDK7) is required for its clearance from promoters (5). In cells, RNAP II is already engaged at promoters of most inactive and inducible genes. After promoter clearance, RNAP II departs from the transcription start site (TSS), but pauses after transcribing 20-100 nucleotide-long transcripts (6,7) by the negative elongation (NELF) (8) and DRB sensitivity inducing factors (DSIF) (9). To release RNAP II for productive elongation, the positive transcription factor b (P-TEFb) is required (10). P-TEFb is composed of CDK9 and C-type cyclins T1 or T2 (CycT1 or CycT2) (11). While CycT1 is expressed ubiquitously, CycT2 is more restricted (12). CDK9 is a Ser/Thr/Pro-directed kinase (i.e. PITALRE) (13), that phosphorylates Ser2 of RNAPII CTD as well as the Spt5 subunit of DSIF and NELF-E subunit of NELF, which releases RNAPII from the paused state (14,15). Thus, P-TEFb is essential for transcriptional elongation.

To maintain the appropriate state of proliferation and growth of cells, the kinase activity of P-TEFb is tightly controlled. In proliferating cells, 7SK small nuclear ribonucleoprotein (7SK snRNP) complex sequesters the majority (50% to 90%) of P-TEFb in an inactive state (16). Besides P-TEFb, 7SK snRNP consists of the abundant 7SK small nuclear RNA (7SK snRNA), hexamethylene bisacetamide (HMBA)-inducible mRNAs 1 and 2 (HEXIM1/2) proteins, La-related protein 7 (LARP7) and methyl phosphate capping enzyme (MePCE), where HEXIM1/2 and 7SK snRNA inhibit the kinase activity of CDK9 (2,17). Various physiological, developmental, and environmental stimuli release P-TEFb from 7SK snRNP and stimulate its kinase activity, thus regulating differentiation, growth and proliferation of cells (18,19). Moreover, various solid tumors, leukemias and lymphomas, tissue inflammation, cardiac hypertrophy (20), depend on the dysregulation of this P-TEFb equilibrium.

While P-TEFb levels are high in proliferating cells, it is absent in quiescent cells such as memory T cells due to the vanishingly low levels of CycT1 in these cells(21–23). CDK9 levels remain high. Chaperone proteins HSP70 and HSP90 maintain its stability (24). Since transcription of viral genes requires P-TEFb, the disappearance of P-TEFb plays a major role in the latency of the human immunodeficiency virus type 1 (HIV-1) (23,25). In these cells, transcripts for CycT1 and CDK9 remain high (21,26), indicating that CycT1 protein levels are regulated post-transcriptionally. Recently, we demonstrated that CycT1 unbound to CDK9 (free CycT1) is rapidly degraded by the proteasome, which occurs in resting T cells as well as in unresponsive T cells, which include anergic and exhausted T cells (27). Whereas, P-TEFb assembly is promoted by the phosphorylation of CycT1 by PKC, its dephosphorylation by protein phosphatase 1 (PP1) causes its dissociation from CDK9 and subsequent degradation (27). To further clarify the mechanism of CycT1 degradation, we first mapped its degron. Next, we identified Siah1 (seven in absentia homolog 1) and Siah2 as E3 ligases that bind to and ubiquitinate CycT1. Finally, inhibiting Siah1/2 increased levels of free CycT1 in resting T cells. Thus, the degradation of free CycT1 by Siah1/2 regulates levels of P-TEFb in quiescent cells.

## MATERIALS AND METHODS

| Key Resources Table |  |  |  |  |
| --- | --- | --- | --- | --- |
| Resource | Designation | Source or reference | Identifiers | Additional information |
| Antibody | CycT1 (E-3) (mouse monoclonal) | Santa Cruz Biotechnology | sc-271348 | WB, 1:200 |
| Antibody | CDK9 (F-6) (mouse monoclonal) | Santa Cruz Biotechnology | sc-376646 | WB, 1:200; IP, 4 µg per test |
| Antibody | CycT1 (D1B6G) (rabbit monoclonal) | Cell Signaling Technology | 81464S | WB, 1:1000 |
| Antibody | CDK9 (EPR22956-37) (rabbit polyclonal) | Abcam | ab239364 | WB, 1:1000; IP, 4 µg per test |
| Antibody | HA (clone HA-7) (mouse monoclonal) | Sigma-Aldrich | H3663 | WB, 1:1000; IP, 4 µg per test |
| Antibody | Flag (clone M2) (mouse monoclonal) | Sigma-Aldrich | F1804 | WB, 1:1000; IP, 4 µg per test |
| Antibody | HA (rabbit monoclonal) | Sigma-Aldrich | H6908 | WB, 1:1000; IP, 4 µg per test |
| Antibody | Flag (rabbit polyclonal) | Sigma-Aldrich | F7425 | WB, 1:1000; IP, 4 µg per test |
| Antibody | Myc-tag (9B11) (mouse monoclonal) | Cell Signaling Technology | 2276S | WB, 1:2000 |
| Antibody | β-Actin (13E5) (rabbit monoclonal) | Cell Signaling Technology | 4970S | WB, 1:5000 |
| Antibody | Siah1 (goat polyclonal) | Abcam | ab2237 | WB, 1:500 |
| Antibody | Siah2 (rabbit polyclonal) | Proteintech | 12651-1-AP | WB, 1:350 |
| Antibody | normal rabbit control IgG | Santa Cruz Biotechnology | sc-2027 | IP, 4 µg per test |
| Antibody | normal mouse control IgG | Santa Cruz Biotechnology | sc-2050 | IP, 4 µg per test |
| Antibody | mouse anti-goat IgG-HRP | Santa Cruz Biotechnology | sc-2354 | WB, 1:5000 |
| Antibody | Amersham ECL Mouse IgG, HRP-linked whole Ab (from sheep) | Cytiva | NA931 | WB, 1:10000 |
| Antibody | Amersham ECL Rabbit IgG, HRP-linked whole Ab (from donkey) | Cytiva | NA934 | WB, 1:10000 |
| Antibody | PE-CF594 mouse anti-human CD69 | BD Biosciences | 562617 | Flow Cytometry, 5 µl per test |
| Antibody | PE-CF594 mouse IgG1 | BD Biosciences | 562292 | Flow Cytometry, 5 µl per test |
| Chemical compound drug | bortezomib | Calbiochem | 179324-69-7 |  |
| Chemical compound drug | MG132 | Sigma-Aldrich | M7449 |  |
| Chemical compound drug | MLN4924 | Calbiochem | 951950-33-7 |  |
| Chemical compound drug | cycloheximide (CHX) | Sigma-Aldrich | C4859 |  |
| Chemical compound drug | N,N'-Hexamethylene bis(acetamide) (HMBA) | Sigma-Aldrich | 224235 |  |

### Plasmids, Antibodies, Chemicals and Proteins

Plasmids containing HA-CycT1 (h:CycT1), CDK9-Flag (CDK9:f) and other mutated or truncated CycT1 cDNAs were constructed by cloning PCR fragments amplified from the coding sequences of CycT1 and CDK9 into pcDNA3.1 with indicated epitope tags. Plasmids containing human Siah1, mutant Siah1C75S and Siah2 cDNAs were kindly provided by Prof. Qiang Zhou (UC Berkeley); plasmids containing mouse Siah2, mutant Siah2RM, PHYL(130) cDNAs and indicated pLKO-shRNAs targeting Siah1 and Siah2, and specific Siah inhibitors (RLS-12, RLS-24, RLS-27 and RLS-96) were kindly provided by Prof. Ze’ev Ronai (Sanford Burnham Prebys). All commercially available antibodies and compounds used in this study are listed in Key Resource Table (above).

### Cell culture

Human Embryonic Kidney (HEK) 293T cells were cultured in Dulbecco’s modified Eagle’s medium (DMEM) (Corning) with 10% fetal bovine serum (FBS) (Sigma Aldrich), Peripheral Blood Mononuclear Cells (PBMCs) were cultured in Roswell Park Memorial Institute (RPMI) 1640 (Corning) with 10% FBS at 37 °C and 5% CO2. Resting CD4+ T cells were purified from bulk PBMCs by using Dynabeads™ Untouched™ Human CD4 T Cells Kit (ThermoFisher Scientific).

### Cell manipulation

Transfection of plasmid DNA was conducted in 293T cells using Lipofectamine 3000 (Life Technology) and X-tremeGENE™ HP DNA Transfection Reagent (Roche) according to the manufacturer’s instructions. Electroporation of plasmid DNA was conducted in resting CD4+ T cells by using Neon™ Transfection system (Invitrogen) through the suggested protocol for PBMC according to the manufacturers’ instructions. 293T cells were treated with 2 µM bortezomib for 8-12 h, 100 µg/ml CHX for 3h, 6h or 9h, and resting CD4+ T cells were treated with 100 nM bortezomib for 12 h, 5 nM phorbol myristate acetate (PMA) (Sigma Aldrich) and 15 ng/ml phytohaemagglutinin (PHA) (Sigma Aldrich) for 24 h before the cell lysis. 293T cells and resting CD4+ T cells were treated with indicated concentrations of Siah inhibitors for 12 h before the cell lysis.

### Co-immunoprecipitation (Co-IP)

293T cells were lysed with chilled RIPA buffer (50 mM Tris-HCl, pH 8.0, 5 mM EDTA, 0.1% SDS, 1.0% Nonidet P-40, 0.5% sodium deoxycholate, 150 mM NaCl) containing protease and phosphatase inhibitors. After brief sonication (level 4, 2 s, once) and centrifugation (21,000×g, 10min), the supernatant was precleared with protein G-Sepharose beads for 2 h. The precleared supernatant was incubated with indicated primary antibodies or control IgG overnight, and incubated with protein G-Sepharose beads for additional 2 h. Beads were washed 5 times with a high-salt RIPA buffer (500 mM NaCl). The co-IP samples and input (1% of whole cell lysates) were boiled in 2× Laemmli sample buffer (Bio-Rad) supplemented with 2-mercaptoethanol (Bio-Rad) and subjected to western blotting (WB) as described previously (55).

### Ubiquitination assays

Two assays (IP and Ni-ATA pulldown) were employed to measure protein ubiquitination in cells transiently expressing His-Ubiquitin. IP was conducted with anti-His antibodies as described above, except with more thorough sonication (level 4, 5s, 4 cycles) before centrifugation. Ni-NTA pull down was conducted by using HIS-Select Nickel Affinity Gels (Sigma). After lysing cell pellet with buffer A (6 M guanidine-HCl, 0.1 M Na_2_HPO_4_/NaH_2_PO_4_, 10 mM imidazole), lysates were briefly sonicated (level 5, 5s, 4 cycles) on ice, and rotated at room temperature for 3 h. Beads were then washed sequentially: once with the buffer A, twice with the buffer B (1.5 M guanidine-HCl, 25 mM Na_2_HPO_4_/NaH_2_PO_4_, 20 mM Tris-Cl pH 6.8, 17.5 mM imidazole), and twice with the buffer TI (25 mM Tris-Cl pH 6.8, 20 mM imidazole). Beads containing ubiquitin-conjugated proteins were then boiled in 2× Laemmli sample buffer containing 2-mercaptoethanol and 200 mM imidazole. IP or pull down samples were subjected to WB as described above.

### Quantification of WBs

WBs were visualized by enhanced chemiluminescence (ECL) (Perkin Elmer) produced by HRP-conjugated secondary antibodies, and chemiluminescent signals were directly captured by LI-COR image analyzer. Band intensities of WBs were quantified using Image Studio software (LI-COR). Relative protein expression in whole cell lysates was calculated by normalizing the indicated proteins with loading control β-actin. Relative protein-protein interactions in IP and co-IP or pulldown were calculated by normalizing the IPed proteins with indicated antibodies targeted proteins or input proteins. Quantification data were presented as fold change over values obtained with control samples.

### RNA extraction and reverse transcription qPCR (RT-qPCR)

Cells were washed twice with chilled PBS, and lysed with the TRIzol reagent (Life Technologies). Total RNA was extracted by phase-separation, and genomic DNA was removed by using the Turbo DNA kit (Life Technologies) following the protocol provided by the manufacturer. The same amount (3 µg) of total RNA was reverse-transcribed into cDNAs by using SuperScript III reverse transcriptase (Life Technologies). Levels of cDNAs corresponding to the indicated mRNAs were measured by qPCR using the SensiFAST SYBR Lo-ROX Kit on the Mx3005p thermalcycler, by normalizing data with the mRNA levels of a house-keeping glyceraldehyde-3-phosphate dehydrogenase (GAPDH).

## RESULTS

### Free CycT1 is highly unstable in cells

Levels of P-TEFb are reduced greatly due to the absence of CycT1 in quiescent and terminally differentiated cells (23,28,29). Nevertheless, mRNA levels of CycT1 and CDK9 remain high in these cells, indicating that post-transcriptional steps are responsible (27,30). Consistent with previous studies, we also confirmed that CycT1 is stabilized by the proteasomal inhibitor bortezomib (100 nM, 24 h) in resting T cells to levels equivalent to those in activated T cells (Figure 1A, upper panels, panel 1, compare lane 2 and 3 to lane 1, ~14.1-fold and ~18.2-fold increase) (27). Levels of CDK9 were not affected by these treatments (Figure 1A, upper panels, panel 2). In addition, mRNA levels of CycT1 were not increased under these different conditions (Figure 1A, middle bar graph). Furthermore, bortezomib did not increase the expression of the activation marker CD69, which is also consistent with previous reports (Figure 1A, bottom column, compare lane 2 to lanes 1 and 3) (31).

**Figure 1.**
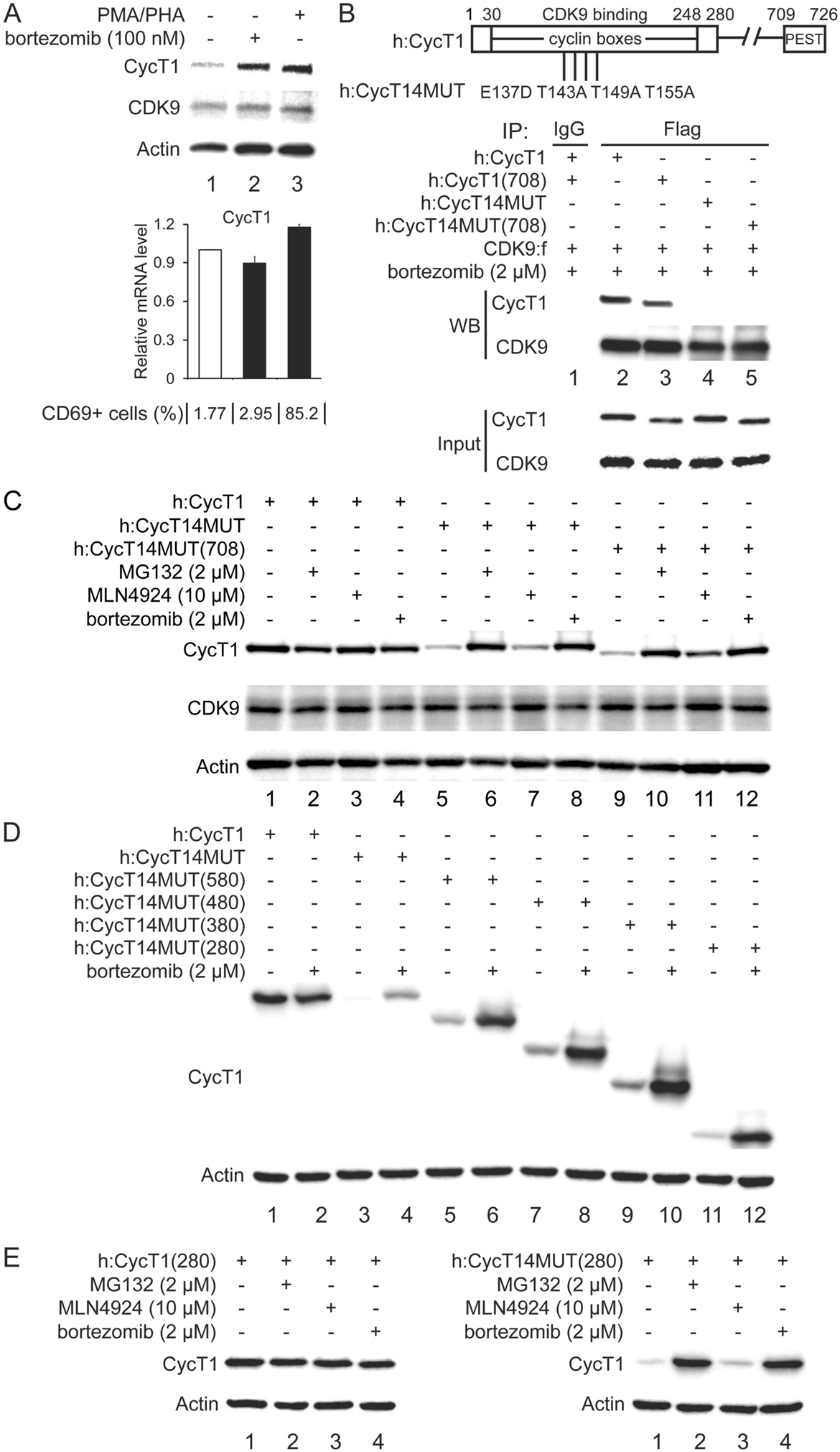
Free CycT1 proteins are unstable but stabilized by proteasomal inhibitors in cells. A. Levels of endogenous CycT1 proteins in resting CD4+ T cells are increased by proteasomal inhibitors without mRNA induction and cell activation. Resting CD4+ T cells were untreated (lane 1) or treated with 100 nM bortezomib (lane 2) or PMA/PHA (lane 3) for 24 h before cell lysis. Levels of CycT1 (panel 1), CDK9 (panel 2), and the loading control actin (panel 3) proteins were detected with indicated antibodies, respectively, by WB. Relative mRNA levels of CycT1 in bortezomib or PMA/PHA treated cells are presented in the middle bar graph as - fold change above levels of untreated cells (white bars, set as ‘1’). Error bars represent standard error of average (n = 3). Percentage of CD69+ cells in CD4+ T cells with indicated treatment are presented in the bottom table by flow cytometry with PE-conjugated anti-CD69 antibodies.
B. Graphic representation of mutant CycT1 proteins that do not interact with CDK9. Top diagram is a schematic presentation of the WT and mutant CycT1 proteins used in this study. WT CycT1 protein contains 726 residues, in which two cyclin boxes are found the N-terminal region (30-248) and a PEST motif is located at its C-terminus (709-726). The mutant CycT1 protein (CycT14MUT) includes Thr143, Thr149, Thr155 and Glu137. WT or indicated mutant CycT1 and CDK9 proteins were expressed in 293T cells treated with bortezomib and co-IPed with CDK9 by anti-Flag antibodies (bottom panels 1 and 2). Bottom panels 3 and 4 indicate input levels of CycT1 and CDK9 proteins.
C. Mutant CycT1 proteins that do not bind to CDK9 are degraded via the proteasome. WT CycT1 or two indicated mutant CycT1 proteins were expressed in 293T cells, which were untreated (lanes 1, 5, and 9) or treated with 2 µM MG132 (lanes 2, 6, and 10), bortezomib (lanes 4, 8, and 12), 10 µM MLN4924 (lanes 3, 7, and 11) before cell lysis. Levels of CycT1 (panel 1), CDK9 (panel 2) and the loading control actin (panel 3) proteins were detected with indicated antibodies, respectively, by WB.
D. C-terminally truncated mutant CycT1 proteins are unstable. WT CycT1 or five indicated mutant CycT1 proteins (CycT14MUT, CycT14MUT(580), CycT14MUT(480), CycT14MUT(380) and CycT14MUT(280)) were expressed in 293T cells in the presence of 2 µM bortezomib (lanes 2, 4, 6, 8, 10 and 12) or not (lanes 1, 3, 5, 7, 9 and 11) before cell lysis. Levels of CycT1 (panel 1) and the loading control actin (panel 2) proteins were detected with indicated antibodies, respectively, by WB.
E. Mutant CycT14MUT(280) proteins are degraded via proteasomal pathways. WT CycT1(280) (left panels) or mutant CycT14MUT(280) (right panels) proteins were expressed in 293T cells, which were untreated (lanes 1) or treated with 2 µM MG132 (lanes 2), bortezomib (lanes 4), 10 µM MLN4924 (lanes 3) before cell lysis. Levels of CycT1 (panels 1) and the loading control actin (panels 2) proteins were detected with indicated antibodies, respectively, by WB.

Recently, we demonstrated that PKCs phosphorylate CycT1 on two threonines (Thr143 and Thr149), which increases its binding to CDK9. Indeed, the stability of CycT1 is determined by its binding to CDK9 (P-TEFb assembly) (27). We sought to clarify the mechanism by which free CycT1 is degraded in cells. Previously, the C-terminal region (amino acid 706-726) of CycT1 was implicated (11). It contains several Pro-, Glu-, Ser-, and Thr-residues, forming a potential PEST motif that has been associated with the degradation of other proteins (32–34). Therefore, we examined if this PEST contributes to the degradation of free CycT1. Also, we employed a mutant CycT1 protein (CycT14MUT, Figure 1B) that does not interact with CDK9. This mutant CycT14MUT protein recapitulates the degradation of free CycT1, which occurs in resting, terminally differentiated, anergic and exhausted cells, in growing, proliferating cells, such as human embryonal kidney (HEK) 293T cell line (27).

First, interactions between WT CycT1 or mutant CycT14MUT proteins in the full length or PEST-truncated versions and CDK9 were analyzed by co-immunoprecipitation (co IP) (Figure 1B). Consistent with our previous studies, the mutant CycT14MUT and CycT14MUT(708) proteins did not interact with CDK9, compared to the WT CycT1 or CycT1(708) proteins (Figure 1B, panel 1, compare lanes 4 and 5 to lanes 2 and 3). Also, the WT CycT1 and truncated mutant CycT1(708) did not disappear after the addition of cycloheximide (CHX) to block protein synthesis (data not presented), indicating that both proteins are equally stable in cells. In contrast, levels of mutant CycT1 proteins (CycT14MUT and CycT14MUT(708)) were significantly decreased (Figure 1C, panel 1, compare lines 5 and 9 to lane 1, ~15.1-fold and ~13.9-fold reduction). However, levels of these proteins were restored by treating cells with proteasomal inhibitors, MG132 or bortezomib, but not with the protein neddylation inhibitor MLN4924 (Figure 1C, panel 1, compare lanes 6 and 8 lane 5 versus lane 7 to lane 5, ~13.5-fold and ~14.3-fold versus ~1.3-fold increase; compare lanes 10 and 12 to lane 7 versus lane 11 to lane 7, ~14.1-fold and ~12.9-fold versus ~1.6-fold increase), indicating that these proteins were degraded by neddylation-independent proteasomal pathways. Levels of the WT CycT1 (Figure 1C, panel1, lanes 1 to 4) and endogenous CDK9 (Figure 1C, panel 2) proteins were not affected by these inhibitors. Thus, consistent with our previous studies, mutant CycT1 proteins that do not interact with CDK9 are highly unstable in cells, which is independent of the C-terminal PEST motif.

To determine the region of CycT1 responsible for its degradation (degron), we constructed several C-terminally truncated WT and mutant CycT14MUT proteins. As presented in Figure 1D, compared to the WT CycT1 (panel 1, lanes 1 and 2), levels of the mutant CycT14MUT and other four C-terminally truncated mutant CycT14MUT proteins were significantly lower in the absence of bortezomib (panel 1, compare lanes 3, 5, 7, 9, and 11 to lane 1, range from ~12.9 to 35.7-fold reduction). They were increased significantly by bortezomib (panel 1, compare lanes 4, 6, 8, 10 and 12 to lanes 3, 5, 7, 9 and 11, range from ~10.7 to 18.7-fold increase). Similar to the full length or PEST-truncated CycT1 proteins, levels of the mutant CycT14MUT(280), but not the WT CycT1(280) proteins were increased by MG132 and bortezomib, but not by MLN4924 (Figure 1E, left panel 1, compare lanes 2 to 4 to lane 1; right panel 1, compare lanes 2 and 4 to lane 1 versus lane 3 to lane 1, ~17.1-fold and ~15.9-fold versus ~1.2-fold increase). We conclude that the degron of CycT1 is located within its N-terminal 280 residues.

### Free CycT1 proteins are highly unstable and ubiquitinated in cells

Next, we examined whether the degradation of the mutant CycT14MUT proteins is mediated by ubiquitination. WT CycT1(280) or mutant CycT14MUT(280) proteins were co-expressed with the His-tagged ubiquitin in 293T cells in the presence of bortezomib. Ubiquitinated proteins were immunoprecipitated (IPed) with anti-His antibodies and ubiquituinated-CycT1 proteins were detected by western blotting (WB) with anti-HA antibodies. Compared to the WT CycT1(280) proteins, the mutant CycT14MUT(280) proteins were heavily polyubiquitinated (Figure 2A, panel 1, compare lane 3 to lanes 1, 2 and 4). Since there are multiple lysine residues that can be ubiquitinated, we further substituted these lysines to alanines alone or in combination (Figure 2B). Indicated mutant CycT1 proteins were then co-expressed with His-Ub in 293T cells in the presence of bortezomib. After the pull-down of ubiquitinated proteins by Ni^2+^-NTA resins, mutant CycT1 proteins were detected by WB with anti-HA antibodies. As presented in Figure 2B, six lysine (lysine 247, 253, 265, 266, 268 and 277) residues, which are in or close to the TRM motif of CycT1, were targeted for ubiquitination (Figure 2B, lower panels, compare lane 7 to lanes 1 to 6). Taken together, we conclude that the free CycT1 is heavily ubiquitinated in the region between positions 247 and 277 and is rapidly degraded by the proteasome.

**Figure 2.**
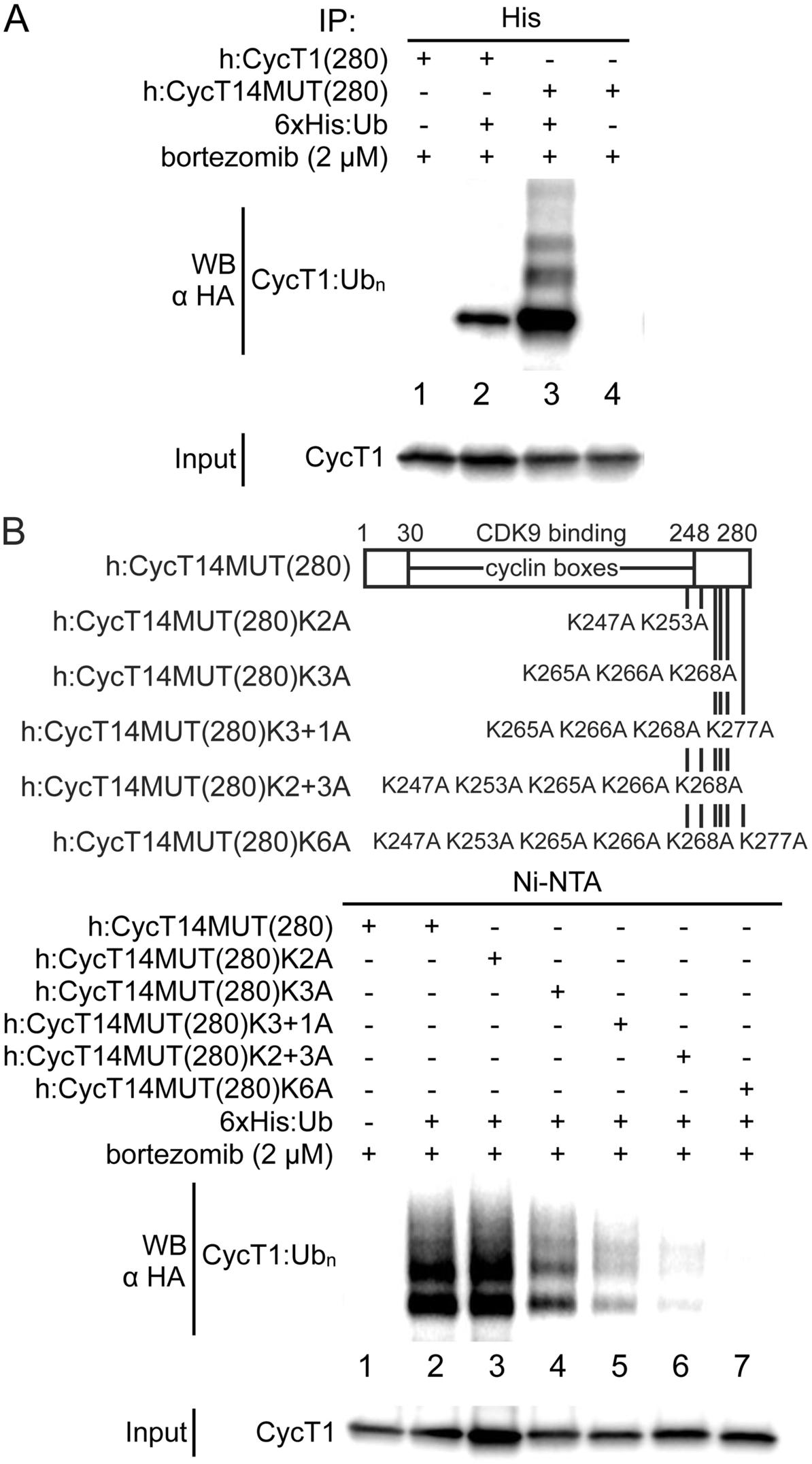
Mapping of the ubiquitination sites of the free CycT1. A. Free CycT1(280) protein is heavily ubiquitinated. WT CycT1(280) or mutant CycT14MUT(280) proteins were co-expressed with (lanes 2 and 3) or without (lanes 1 and 4) His-epitope tagged ubiquitin in 293T cells, followed by treatment with 2 µM bortezomib. Cell lysates were subjected to IPs with anti-His antibodies, and CycT1 proteins were detected with anti-HA antibodies (upper panel). Bottom panel presents input levels of CycT1 proteins.
B. Mapping of ubiquitination sites in the free CycT1(280) protein. Top diagram is a schematic presentation of mutant CycT14MUT(280) proteins and its indicated lysine residues as potential ubiquitination sites. Mutant CycT1(280) proteins contain lysine to alanine (K to A) substitutions at residues 247, 253, 265, 266, 268 and 277. Six indicated mutant CycT14MUT(280) and His-epitope tagged ubiquitin proteins were co-expressed in 293T cells treated with 2 µM bortezomib. Ni-NTA pull down assays were conducted to isolate ubiquitinated CycT1 proteins detected with anti-HA antibodies (upper panel). Bottom panel presents input levels of CycT1 proteins.

### Siah1 and Siah2 preferentially bind to the free CycT1

Since free CycT1 is subjected to ubiquitination and proteasomal degradation, identification of potential E3 ligase(s) responsible for CycT1 degradation is essential for understanding this process. Previous proteomic analysis with P-TEFb revealed that the E3 ligase FBXO11 interacts with the complex (35). Also two other E3 ligases (Grail and Cbl-b) were reported to be up-regulated or enhanced in unresponsive T cells (36,37) where P-TEFb is absent. In our hands, the ectopic expression of FBXO11 or Grail proteins did not decrease levels of the mutant CycT14MUT protein (data not presented). In our previous study, levels of CycT1 were significantly decreased in murine anergic T cells (27), where the depletion of Cbl-b did not affect levels of CycT1 (data not presented). Thus, none of these E3 ligases are involved. Nevertheless, two previous publications reported interactions between AFF1, a component of the Super Elongation Complex (SEC), and the RING domain-containing E3 ubiquitin ligases Siah1 and Siah2 from a yeast two-hybrid screen (38,39). Subsequently, Zhou and colleagues demonstrated that only Siah1 targets the free ELL2 proteins of SEC for ubiquitination-mediated degradation in Hela cells (40,41). Since AFF1, ELL2 and P-TEFb are all components of the SEC complex, we examined whether Siah1 and/or Siah2 also target the free CycT1 in a manner similar to ELL2.

First, we examined if the free CycT1 interacts with Siah1 or Siah2 by co-IP. Since WT Siah1 is reported to be highly unstable because of its self-mediated ubiquitination and degradation (42), its RING domain negative mutant (dominant negative (DN)) Siah1C75S, possessing the same ability to bind to its targets, was used for co-IP. After co-expression of the DN Siah1C75S and WT CycT1 or mutant CycT14MUT proteins in 293T cells in the presence of bortezomib, co-IPs were performed with anti-HA (targeting CycT1) or anti-Flag (targeting Siah1) antibodies. As presented in Figure 3A, a significantly larger amount of the DN Siah1C75S protein was co-IPed with the mutant CycT14MUT protein than with the WT CycT1 protein (panels 1 and 2, compare lane 3 to lane 2, ~5.7-fold increase). Similarly, a larger amount of the mutant CycT14MUT protein compared with the WT CycT1 protein was co-IPed with the DN Siah1C75S protein (Figure 3A, panels 5 and 6, compare lane 3 to lane 2, ~4.3-fold increase). Similar co-IPs were performed by using the WT Siah2 and WT CycT1 or mutant CycT14MUT proteins. Although the WT Siah2 protein also causes self-ubiquitination, its steady-state levels are much higher.Therefore, the WT Siah2 was employed for the co-IP experiments. As presented in Figure 3B, a similar preference of Siah2 interacting with the mutant CycT14MUT over WT CycT1 proteins was observed (panels 1 and 2 and panels 5 and 6, compare lanes 3 to lanes 2, ~8.3-fold and ~10.5-fold increase). Similar co-IPs with a series of C-terminally truncated CycT1 proteins indicated that a region in the N-terminal 280 residues of CycT1 interacts with Siah1 or Siah2 (data not presented). Both the DN Siah1C75S and Siah2 also preferentially interact with the C-terminally truncated mutant CycT14MUT(280) protein, but not with the WT CycT1(280) protein (Figures 3C and 3D, panels 1 and 2, compare lanes 3 to lanes 2, ~5.1-fold and ~8.9-fold increase; panels 5 and 6, compare lanes 3 to lanes 2, ~3.2-fold and ~4.1-fold increase). Thus, similar to the situation with free ELL2, free CycT1 interacts with both Siah1 and Siah2.

**Figure 3.**
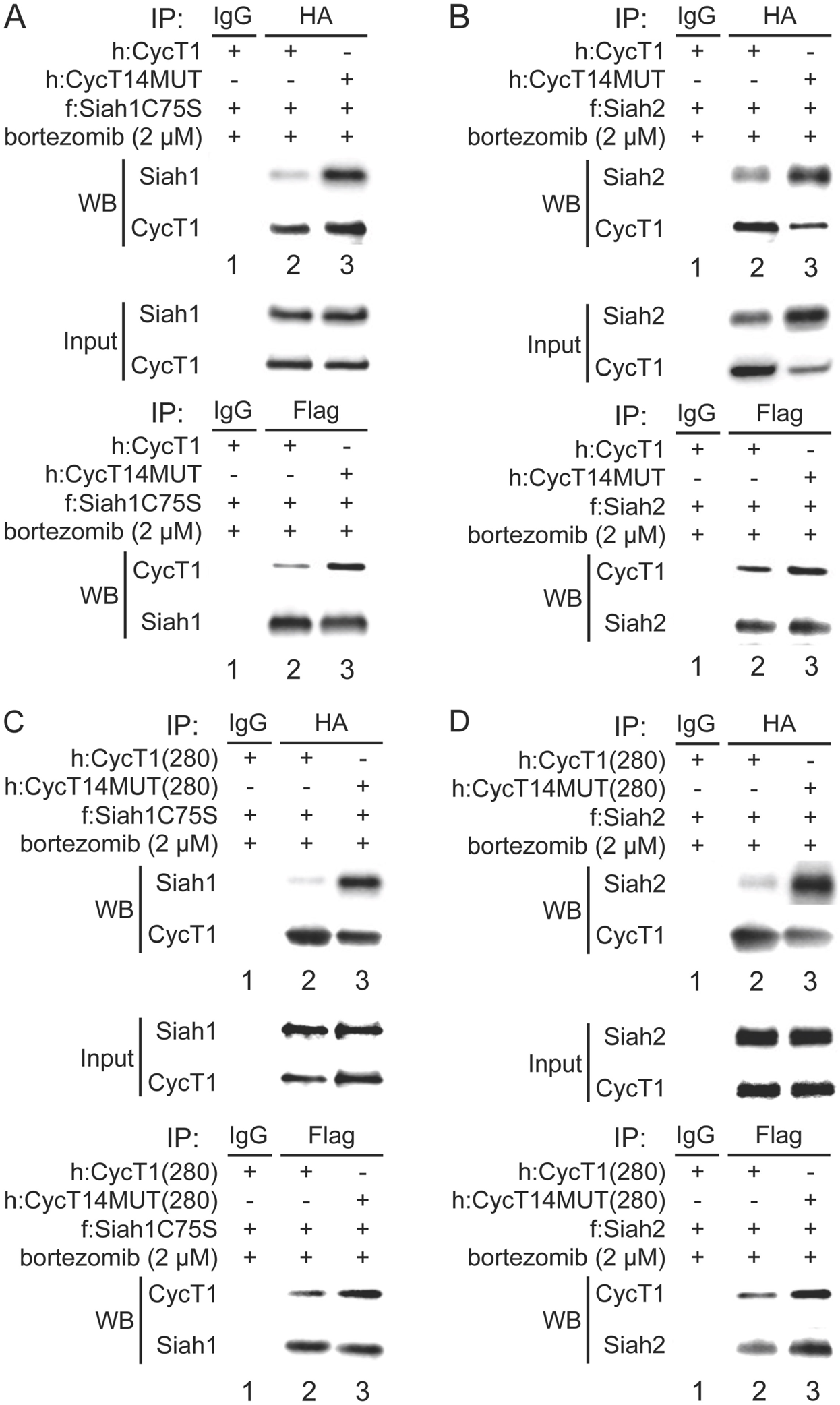
Siah1 and Siah2 preferentially interact with free CycT1. A. Siah1 preferentially interacts with free CycT1 protein. DN Siah1C75S proteins were co-expressed with WT CycT1 or mutant CycT14MUT proteins in the presence of bortezomib in 293T cells, followed by co-IPs with CycT1 (anti-HA antibodies) (panels 1 and 2), or with Siah1 (anti-Flag antibodies) (panels 5 and 6). Panels 3 and 4 indicate input levels of CycT1 and Siah1 proteins.
B. Siah1 preferentially interacts with free CycT1(280) protein. DN Siah1C75S proteins were co-expressed with WT CycT1(280) or mutant CycT14MUT(280) proteins in the presence of bortezomib in 293T cells, followed by co-IPs with CycT1 (anti-HA antibodies) (panels 1 and 2), or with Siah1 (anti-Flag antibodies) (panels 5 and 6). Panels 3 and 4 indicate input levels of CycT1 and Siah1 proteins.
C. Siah2 preferentially interacts with free CycT1 protein. WT Siah2 proteins were co-expressed with WT CycT1 or mutant CycT14MUT proteins in the presence of bortezomib in 293T cells, followed by co-IPs with CycT1 (anti-HA antibodies) (panels 1 and 2), or with Siah2 (anti-Flag antibodies) (panels 5 and 6). Panels 3 and 4 indicate input levels of CycT1 and Siah2 proteins.
D. Siah2 preferentially interacts with free CycT1(280) protein. WT Siah2 proteins were co-expressed with WT CycT1(280) or mutant CycT14MUT(280) proteins in the presence of bortezomib in 293T cells, followed by co-IPs with CycT1 (anti-HA antibodies) (panels 1 and 2), or with Siah2 (anti-Flag antibodies) (panels 5 and 6). Panels 3 and 4 indicate input levels of CycT1 and Siah2 proteins.

### Identification of a degron motif in CycT1, which is targeted by Siah1 and Siah2

Results described above indicated that the degron of CycT1 is located within the N-terminal 280 residues. To exclude the possibility that CycT1 proteins possess two or more motifs within its N-or C-termini targeted by Siah1/2, co-IPs were performed in cells co-expressing the mutant CycT14MUT(580) or CycT1(430-726) proteins with the DN Siah1C75S or Siah2 proteins. No interactions were observed between the mutant CycT1(430-726) protein and the DN Siah1C75S or Siah2 proteins, compared to significant interactions between the mutant CycT14MUT(580) and both Siah proteins (data not presented). Next, we further mapped the minimum region in CycT1 that interacts with Siah1/2 by examining a series of N-terminal truncated mutant proteins by co-IP. As presented in Figure 4A, whereas the mutant CycT1(260-580) protein did not interact (panels 1 and 2, lane 5), two N-terminally truncated mutant CycT14MUT(100-580) and CycT1(210-580) proteins as well as the mutant CycT14MUT(580) protein strongly interacted with the DN Siah1C75S protein (panels 1 and 2, lanes 2 to 4). Similarly, whereas all other three mutant CycT14MUT proteins interacted strongly with Siah2, the mutant CycT1(260-580) protein did not (Figure 4B, panels 1 and 2, compare lane 5 to lanes 2 to 4). Furthermore, to narrow down the region of CycT1 targeted by Siah1/2, two additional C-terminal truncated mutant CycT14MUT(260) and CycT14MUT(210) proteins, based on the mutant CycT14MUT(280) protein, were constructed and examined. Consistent with the N-terminal truncated mutant proteins, whereas the mutant CycT14MUT(210) protein did not interact with the DN Siah1C75S and Siah2 proteins, mutant CycT14MUT(280) and CycT14MUT(260) proteins did (data not presented). Thus CycT1 from positions 210 to 260 binds to Siah1/2.

**Figure 4.**
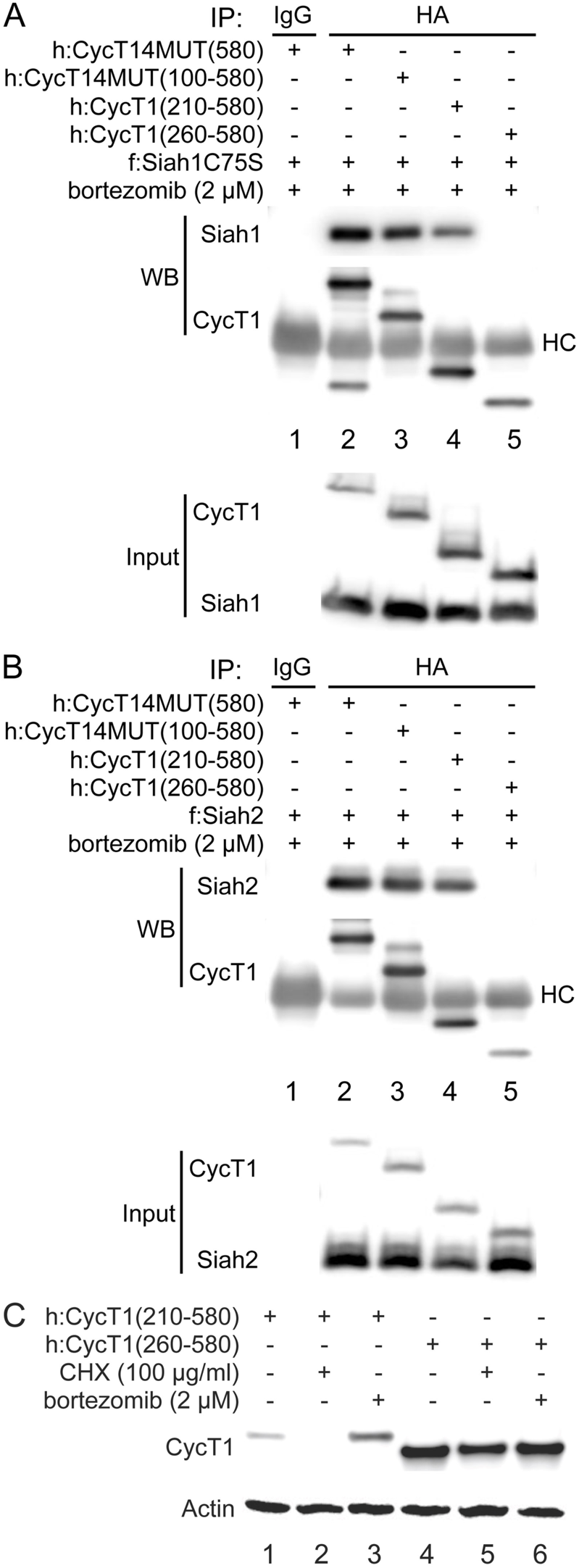
Mapping the region of CycT1 targeted by Siah1/2. A. Mapping the region of CycT1 that interacts with Siah1. DN Siah1C75S proteins were co-expressed with indicated mutant CycT1 proteins in the presence of bortezomib in 293T cells, followed by co-IPs with CycT1 (anti-HA antibodies) (panels 1 and 2). Panels 3 and 4 indicate input levels of CycT1 and Siah1 proteins.
B. Mapping the region of CycT1 that interacts with Siah2. WT Siah2 proteins were co-expressed with indicated mutant CycT1 proteins in the presence of bortezomib in 293T cells, followed by co-IPs with CycT1 (anti-HA antibodies) (panels 1 and 2). Panels 3 and 4 indicate input levels of CycT1 and Siah2 proteins.
C. CycT1 without the degron is stable. Two indicated truncated CycT1 proteins were expressed in 293T cells, which were untreated (lanes 1 and 4) or treated with 100 µg/ml CHX (lanes 2 and 5) or 2 µM bortezomib (lanes 3 and 6) before cell lysis. Levels of CycT1 (panel 1) and the loading control actin (panel 2) proteins were detected with indicated antibodies, respectively, by WB.

Next, to measure the stability of these proteins in cells, the mutant CycT1(210-580) and CycT1(260-580) proteins were expressed in 293T cells in the presence of CHX or bortezomib, followed by WB. As presented in Figure 4C, similar to the mutant CycT14MUT protein, levels of the mutant CycT1(210-580) protein were severely decreased in the presence of CHX, and significantly increased by bortezomib (panel 1, compare lane 2 and 3 to lane 1, ~31.7-fold reduction and ~5.9-fold increase). In contrast, levels of the mutant CycT1(260-580) protein were not affected by CHX or bortezomib (Figure 4C, panel 1, compare lane 6 or lane 5 to lane 3). Taken together, sequences from positions 210 to 260 of CycT1 are targeted by Siah1/2 for binding and degradation. Thus, the removal of this region stabilizes CycT1.

### Siah1/2 degrade the free CycT1

Results presented above indicate that Siah1 and Siah2 are E3 ligases that promote the degradation of free CycT1. To further verify this, we first ectopically expressed Siah1 or Siah2 with the WT or mutant CycT14MUT proteins and their truncated versions (CycT1(280)) without bortezomib in 293T cells, and measured the expression of these proteins by WB. Levels of the WT CycT1 and WT CycT1(280) proteins were minimally affected by co-expression of Siah1 or Siah2, exhibiting only a slight decrease in the presence of the highest levels of Siah1 or Siah2 proteins (data not presented). In contrast, levels of the mutant CycT14MUT and CycT14MUT(280) proteins were significantly decreased by the co-expression of Siah1 and Siah2 in a dose dependent manner (data not presented). The ectopic expression of Siah2 exhibited a greater decrease of mutant CycT1 proteins (data not presented). Notably, high levels of ectopic Siah1/2 did not decrease levels of endogenous bound CycT1 (data not presented).

In addition, we examined if the ectopic expression of mutant Siah1 or Siah2 proteins can rescue the expression of free CycT1 by competing with endogenous Siah1 and Siah2 proteins. As with other RING domain E3 ligases, mutations of catalytic sequences within the RING domain abolish the activity of Siah proteins without reducing their capacity to bind to target proteins (42). Thus, by competing with endogenous Siah proteins, RING domain negative mutant Siah proteins block their degradation (43). Human Siah2 proteins share 85.7% amino acid homology between species, 69.8% identity with the human Siah1 protein, and sequence deviations mainly occur in their N-termini (40,44). Only seven residues are different between human and mouse Siah2 proteins: six are located in the region before the RING domain and one is in the last residue. Activities of these Siah2 proteins from two species are indistinguishable (42). We also observed that mouse Siah2 proteins, like human Siah2 proteins, significantly decrease levels of the mutant CycT14MUT proteins (data not presented). Two RING domain negative mutants, the human DN Siah1C75S and mouse mutant Siah2RM proteins as well as their WT counterparts as controls, were co-expressed with the WT or mutant CycT14MUT proteins in 293T cells. As presented in Figure 5A, WT Siah1 and Siah2 proteins preferentially depleted the mutant CycT14MUT protein (left versus right panels 1 and 2, compare lanes 2 and 3 to lanes 1, ~1.3-fold and ~1.9-fold reduction versus ~7.3-fold and ~11.9-fold reduction). In contrast, RING domain negative Siah mutants, the DN Siah1C75S and mutant Siah2RM proteins stabilized the mutant CycT14MUT protein, but not the WT CycT1 protein (Figure 5A, left versus right panels 1 and 2, compare lanes 4 and 5 to lanes 1, ~1.1-fold and ~1.3-fold increase versus ~9.3-fold and ~6.9-fold increase). Similar results were obtained with the truncated WT CycT1(280) or mutant CycT14MUT(280) and WT or mutant Siah1/2 proteins (Figure 5B, left versus right panels 1 and 2, compare lanes 2 and 3 to lanes 1, ~1.1-fold and ~1.4-fold reduction versus ~5.3-fold and ~9.7-fold reduction; compare lanes 4 and 5 to lanes 1, ~1.6-fold and ~1.4-fold increase versus ~13.3-fold and ~8.9-fold increase). Thus, whereas Siah1/2 preferentially target the free CycT1 for degradation, their RING domain mutant Siah1/2 proteins counteract this effect and stabilize the protein.

**Figure 5.**
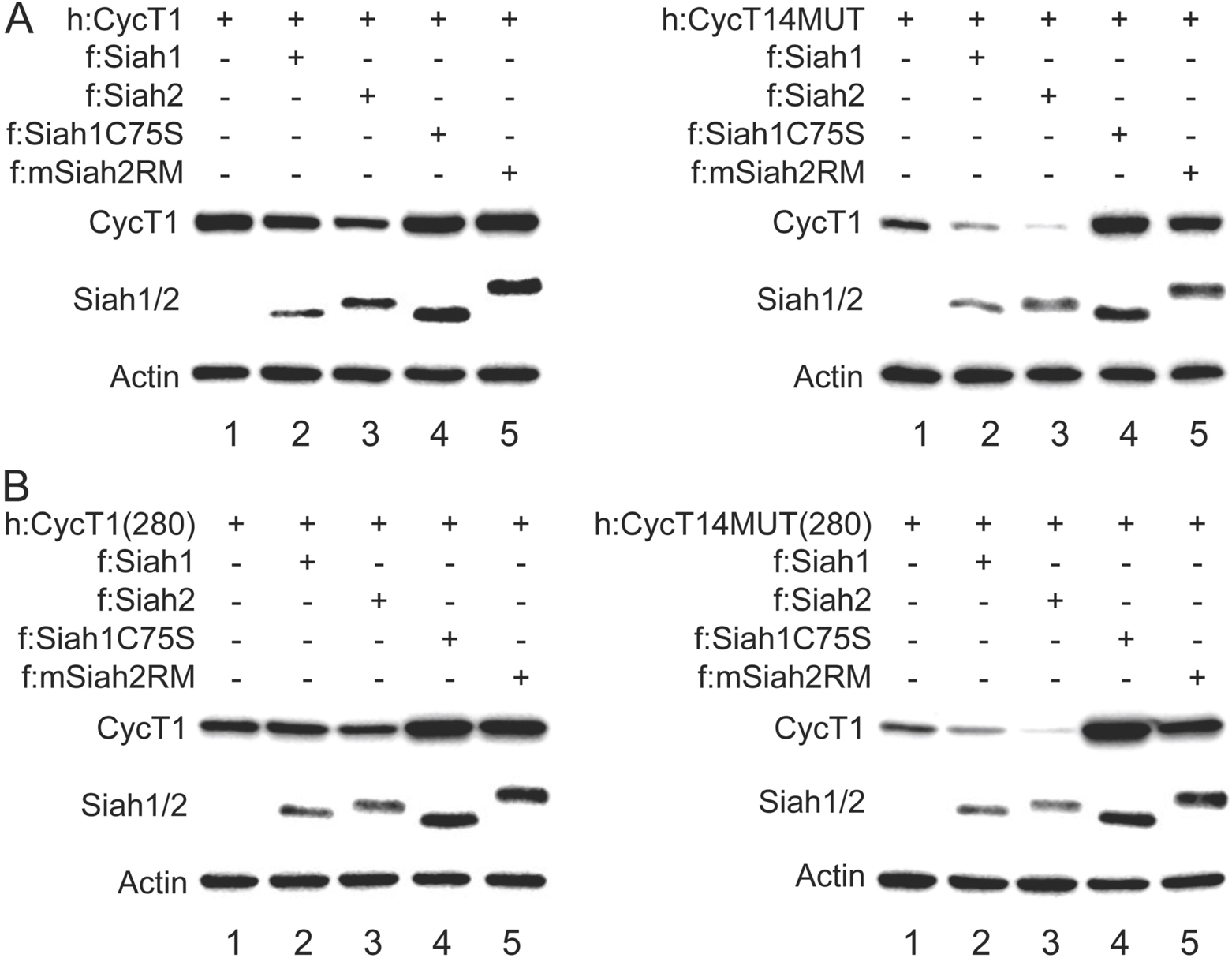
Siah1 and Siah2 preferentially degrade free CycT1, while their dominant negative mutants reverse such degradation. A. Siah1/2 preferentially degrade the free CycT1, while dominant negative mutant Siah1/2 increase its level. WT CycT1 (left panels) or mutant CycT14MUT (right panels) proteins were expressed without (lanes 1) or with WT Siah1 (lanes 2), Siah2 (lanes 3), DN Siah1C75S (lanes 4) or Siah2RM (lanes 5) proteins in 293T cells. Levels of CycT1 (panels 1), Siah1/2 (panels 2) and the loading control actin (panels 3) proteins were detected with indicated antibodies, respectively, by WB.
B. Siah1/2 preferentially degrade free CycT1(280), while dominant negative mutant Siah1/2 increase its level. WT CycT1(280) (left panels) or mutant CycT14MUT(280) (right panels) proteins were expressed without (lanes 1) or with WT Siah1 (lanes 2), Siah2 (lanes 3), DN Siah1C75S (lanes 4) or Siah2RM (lanes 5) proteins in 293T cells. Levels of CycT1 (panels 1), Siah1/2 (panels 2) and the loading control actin (panels 3) proteins were detected with indicated antibodies, respectively, by WB.

### Inhibition or depletion of Siah1/2 stabilize the free CycT1

We demonstrated previously that a drosophila protein phyllopod (PHYL) or its truncated version PHYL(130) are highly potent and specific inhibitors of Siah1/2 (45). They bind and inhibit the substrate-binding domain (SBD) of Siah1/2 (46,47). Therefore, we examined if exogenous PHYL proteins can prevent the degradation of free CycT1. PHYL(130) peptide was co-expressed with the WT CycT1 or mutant CycT14MUT proteins. As presented in Figure 6A, levels of WT CycT1 proteins were not affected by PHYL(130) (left panels, compare lane 3 to lanes 1 and 2). In contrast, levels of the mutant CycT14MUT protein were significantly increased by PHYL(130), to those equivalent to bortezomib treatment for 8 h (Figure 6A, right panels, compare lanes 2 and 3 to lane 1, ~15.9-fold and ~13.5-fold increase). These results further confirm that inhibition of endogenous Siah proteins reverses the degradation of free CycT1 in 293T cells.

**Figure 6.**
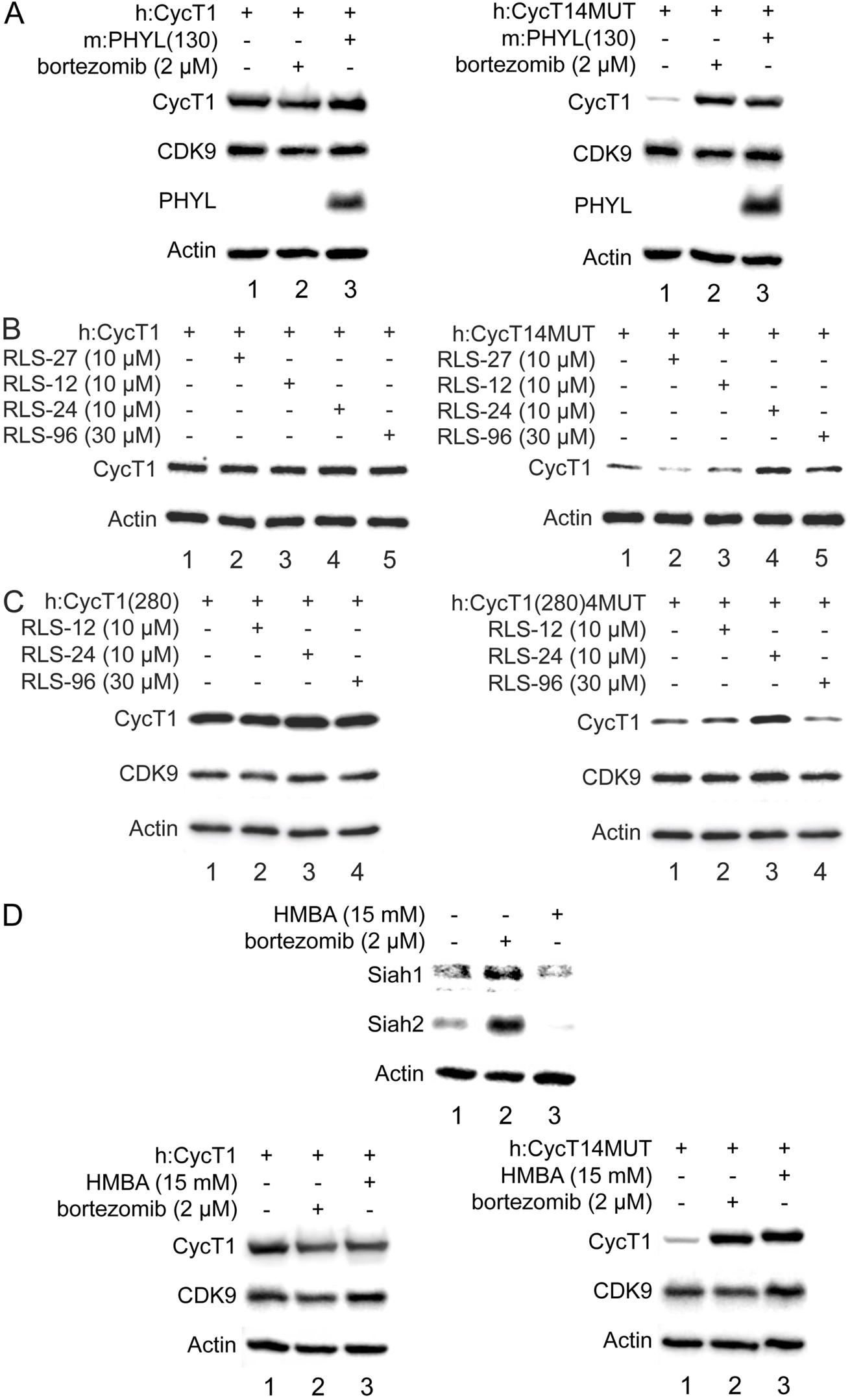
Inhibition or depletion of Siah1/2 increases levels of free CycT1. A. Ectopic expression of a Siah inhibitory peptide PHYL(130) increases levels of free CycT1. WT CycT1 (left panels) or mutant CycT14MUT (right panels) proteins were expressed in the absence (lanes 1) or presence (lanes 2) of bortezomib, or co-expressed with PHYL(130) without bortezomib (lanes 3) in 293T cells. Levels of CycT1 (panels 1), CDK9 (panels 2), PHYL (panels 3) and the loading control actin (panels 4) proteins were detected with indicated antibodies, respectively, by WB.
B. Inhibitors of Siah1/2 increase levels of free CycT1. WT CycT1 (left panels) or mutant CycT14MUT (right panels) proteins were expressed in 293T cells. Cells were untreated (lanes 1) or treated with indicated Siah1/2 inhibitors (lanes 2 to 5). Levels of CycT1 (panels 1) and the loading control actin (panels 2) proteins were detected with indicated antibodies, respectively, by WB.
C. Inhibitors of Siah1/2 increase levels of free CycT1(280). WT CycT1(280) (left panels) or mutant CycT14MUT(280) (right panels) proteins were expressed with in 293T cells. Cells were untreated (lanes 1) or treated with indicated Siah1/2 inhibitors (lanes 2 to 4). Levels of CycT1 (panels 1), CDK9 (panels 2) and the loading control actin (panels 3) proteins were detected with indicated antibodies, respectively, by WB.
D. Depletion of Siah1/2 by HMBA increases levels of free CycT1. 293T cells (top panels) treated without (lane 1) or with 2 µM bortezomib for 12 h (lane 2) or 15mM HMBA for 24 h (lane 3). Levels of Siah1 (panel 1), Siah2 (panel 2) and the loading control actin (panel 3) proteins were detected with indicated antibodies, respectively, by WB. WT CycT1 (bottom left panels) or mutant CycT14MUT (bottom right panels) proteins were expressed in 293T cells with the same treatment as above. Levels of CycT1 (panels 1), CDK9 (panels 2) and the loading control actin (panels 3) proteins were detected with indicated antibodies, respectively, by WB.

In addition, we developed several chemical inhibitors of Siah1/2 (48). These Siah inhibitors were examined for the inhibition of free CycT1 degradation by Siah1/2 in 293T cells. As presented in Figure 6B, indicated amounts of four different compounds (RLS-12, 24, 27, 96) were added to cells for 12 h before subjecting them to WB. Among these four different compounds, no inhibitory effects were detected with RLS-27 (Figure 6B, left and right panels, compare lanes 2 to lanes 1), which was used as the negative control. Effects of the other three compounds on Siah1/2 are varied depending on cell types (48). While levels of WT CycT1 proteins were not affected by any of these inhibitors (Figure 6B, left panels), those of the mutant CycT14MUT protein were significantly increased by RLS-24, and increased to lesser extent by RLS-12 and RLS-96 (Figure 6B, right panel 1, compare lane 4 to lane 1 versus lanes 3 and 5 to lane 1, ~6.3-fold increase versus ~1.6-fold and ~2.7-fold increase). Similarly, these three inhibitors, RLS-12, 24 and 96, also increased levels of the mutant CycT14MUT(280) protein (Figure 6C, right panel 1, compare lane 3 to lane 1 versus lanes 2 and 4 to lane 1, ~4.7-fold increase versus ~1.4-fold and ~1.3-fold increase), but did not affect levels of the WT CycT1(280) protein (Figure 6C, left panels). Thus, chemical inhibition of Siah1/2 also increases levels of free CycT1.

Previously, Zhou and colleagues demonstrated that levels of endogenous Siah1 proteins and its mRNA are severely diminished by a potent differentiation-inducer HMBA in Hela cells, resulting in stabilizing the free ELL2 without increasing its mRNA levels (40). We also observed that levels of Siah1 and Siah2 were greatly decreased by HMBA in 293T cells (Figure 6D, top panels 1 and 2, compare lanes 3 to lanes 1, ~3.9-fold and ~7.9-fold reduction). In contrast, levels of Siah1/2 proteins were increased by bortezomib (Figure 6D, top panels 1 and 2, compare lanes 2 to lanes 1, ~5.9-fold and ~6.9-fold increase), indicating that these proteins are degraded in the presence of HMBA. Also, consistent with previous studies, mRNA levels of Siah1 and Siah2 were decreased by HMBA treatment (data not presented). Thus, transcriptional and post-transcriptional mechanisms are involved in the HMBA-mediated depletion of Siah1/2. Next, we measured levels of the ectopically expressed WT CycT1 or mutant CycT14MUT proteins in the presence of HMBA. As presented in Figure 6D, levels of mutant CycT14MUT proteins were significantly increased by HMBA to those equivalent to bortezomib. In contrast, levels of the WT CycT1 were not affected by HMBA or bortezomib (Figure 6D, bottom left panel 1, compare lanes 2 and 3 to lane 1). Levels of endogenous CDK9 were not affected by these treatments (Figure 6D, bottom panels 2). We conclude that depletion of Siah1/2 by other means also restores revels of free CycT1, further supporting the notion that they are the E3 ligases that degrade this protein.

Next, we targeted the endogenous Siah2 protein by shRNA targeting Siah2 into 293T cells. 48 h after transfection of specific shRNA pool or scrambled RNA as a control, the WT or mutant CycT14MUT proteins were expressed by transient transfection for another 48 h before WB. With significantly decreased levels of Siah2 by shRNA (Figure 7A, panel 1, compare lane 2 to lane 1, ~6.7-fold reduction), levels of the mutant CycT14MUT but not WT CycT1 proteins were greatly increased (Figure 7A, panel 3 versus panel 5, compare lanes 2 to lanes 1, ~1.2-fold increase versus ~5.9-fold increase). Similarly, levels of the mutant CycT14MUT(280) but not WT CycT1(280) proteins were also increased greatly in Siah2 depleted cells (data not presented). Since Siah1 is rather unstable in these cells, this finding confirms the involvement of Siah1/2 in the degradation of free CycT1.

**Figure 7.**
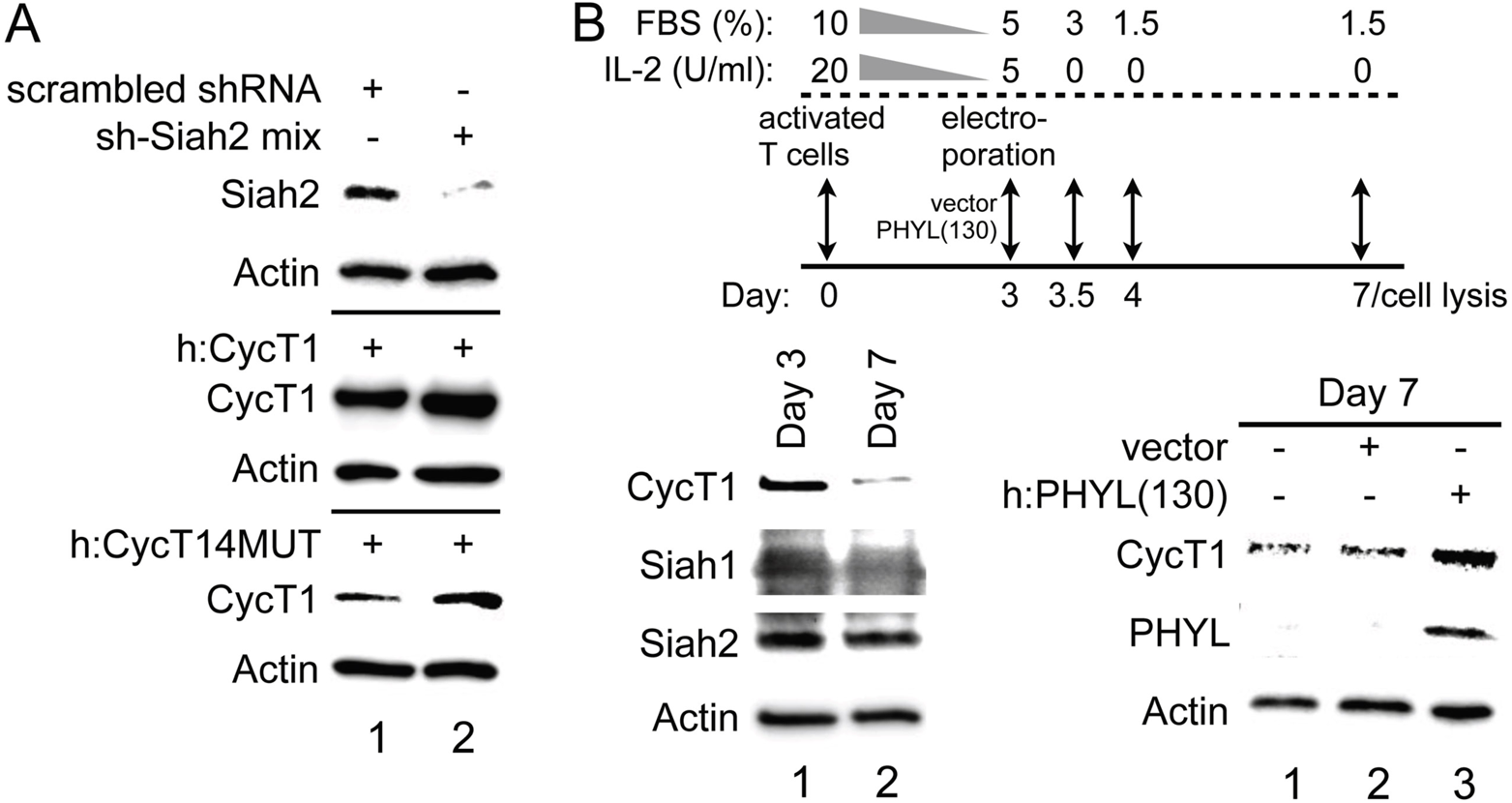
Knockdown of Siah2 increases levels of free CycT1 in 293T cells and inhibition of Siah1/2 stabilizes CycT1 in primary T cells. A. Knockdown of Siah2 stabilizes free CycT1 in 293T cells. 293T cells were transfected with scrambled shRNA (lane 1) or mixed shRNA targeting Siah2 (lane 2). 48 h after transfection, WT CycT1 or mutant CycT14MUT proteins were expressed by transient transfection for another 48 h before cell lysis. Levels of Siah2 (panel 1), CycT1 (panels 3 and 5) and the loading control actin (panels 2, 4 and 6) proteins were detected with indicated antibodies, respectively, by WB.
B. Ectopic expression of PHYL(130) stabilizes CycT1 in primary CD4+ T cells. Top diagram presents the experimental scheme. Primary T cells collected at days3 and 7 (bottom right panels, lanes 1 and 2) were lysed and subjected to WB. Levels of CycT1 (panel 1), Siah1 (panel 2), Siah2 (panel 3) and the loading control actin (panel 4) proteins were detected with indicated antibodies, respectively. Primary T cells without electroporation (right panels, lane 1) and those electroporated with the empty plasmid vector (right panels, lane 2) or plasmids containing PHYL(130) fragments (right panels, lane 3) were collected at day 7, and cell lysates were subjected to WB. Levels of CycT1 (panel 1), PHYL (panel 2) and the loading control actin (panel 3) proteins were detected with indicated antibodies, respectively.

### Inhibition of Siah1/2 in resting primary cells also rescues the expression of free CycT1

Finally, we examined if Siah1/2 are involved in free CycT1 protein degradation in primary quiescent cells. Extending the findings of previous studies, we first confirmed that proteasomal inhibitors increase levels of CycT1 in resting T cells (Figure 1A) (29). However, RLS-24 has been demonstrated to possess anti-proliferative and pro-apoptotic effects in different cells through altering Caspase 3, Bcl2 and Bax activities (48,49). Here, we also observed increasing cytotoxicity after 12 h treatment with RLS-24 in resting T cells, which lead the decrease most proteins in cells. To use PHYL(130), because of transfection efficiency, we had to first activate cells and then bring them to the resting state by withdrawing IL2. As presented in Figure 7B, we electroporated plasmids with PHYL(130) coding sequences into the primary T cells at day 3 after activation (cells were maintained at 5 U/ml IL-2 and 5% FBS). After removal of IL-2, these cells were maintained in the low FBS (1.5%) containing medium for another four days and were subjected to WB. In these cells, levels of CycT1 decreased significantly after four days (Figure 7B, bottom left panel 1, compare lane 2 to lane 1, ~9.5-fold reduction). Of interest, levels of Siah1 and Siah2 did not change (Figure 7B, bottom left panels 2 and 3, compare lanes 2 to lanes 1). In contrast, at day 7, levels of CycT1 in primary T cells with exogenous PHYL(130) were much higher than in control cells (Figure 7B, bottom right panel 1, compare lane 3 to lane 1 versus lane 2 to lane 1, ~7.9-fold increase versus ~1.2-fold increase). Taken together, we conclude that inhibition or depletion of Siah1/2 proteins increases levels of free CycT1 in proliferating and resting cells.

## DISCUSSION

We demonstrated previously that P-TEFb assembly is mediated by reversible phosphorylation of CycT1, and that the free dissociated CycT1 is subjected to rapid degradation (27). In this study, we identified the detailed mechanism of this degradation. First, ubiquitination and degradation of mutant CycT1 proteins unbound to CDK9 requires its N-terminal region (1-280), but not the C-terminal PEST motif. Free CycT1 is ubiquitinated at six lysines near the TRM region. E3 ligases Siah1 and Siah2 target these N terminal sequences (positions 210 to 260) of free CycT1 for binding and degradation. Inhibition of Siah1/2 rescues the expression of free CycT1 in proliferating as well as resting primary cells. We conclude that Siah1/2 are the E3 ligases that bind to and degrade the free dissociated CycT1 protein.

Previous studies indicated that FBXO11 is associated with P-TEFb (35). However, FBXO11 and other two up-regulated E3 ligases (GRAIL and Cbl-b) are not involved in free CycT1 degradation. Zhou and colleagues previously demonstrated that Siah1 is involved in the degradation of ELL2, which together with P-TEFb, is a critical component of the Super Elongation Complex (SEC) (40). These studies prompted us to examine if Siah1/2 are also involved in the degradation of the free CycT1. Numerous structural and functional studies presented here support this new finding. Together, they explain why the SEC is so heavily impacted by Siah1/2, especially in resting and terminally differentiated/exhausted cells. Our binding and ubiquitination studies revealed that sequences near the C-terminus of the cyclin boxes and TRM in CycT1 are involved. Interestingly, we could find no role for the putative PEST at the extreme C-terminus of CycT1. This PEST might be more functional in different cells or under different conditions, possibly even in the context of the fully assembled P-TEFb. Consistent with the previous data that RLS-24 is the most potent Siah inhibitor in cancer cells (48), RLS-24 led to the most effective rescue of the free CycT1 degradation in 293T cells. All three effective inhibitors were too toxic to primary resting T cells. Thus, we could not examine their effects in resting lymphocytes. Rather PHYL(130) confirmed our findings in these cells. Finally, because Siah1 was already unstable in 293T cells, we only used shRNA against Siah2 to decrease effects of both proteins. Again, because the genetic inactivation of Siah1/2 is lethal (50,51), we could not knock out Siah1/2 in our cells. Rather, highly specific Siah1/2 inhibitory peptide PHYL(130) and the addition of HMBA achieved the same goal and again rescued the expression of the free CycT1. Thus, Siah1/2 not only bind to but degrade free, dissociated CycT1 in cells.

Together with our previous study, we proposed a detailed mechanisms how P-TEFb is regulated in resting and activated cells. In proliferating cells, P-TEFb assembly is promoted by the phosphorylation of CycT1 by PKC, which stabilizes CycT1 by preventing the access of Siah1/2. In resting cells, CycT1 is dephosphorylated by PP1 and dissociates from CDK9, and this free CycT1 is ubiquitinated and degraded by Siah1/2. This situation appears uncannily reminiscent of cell cycle cyclins/CDKs that are also regulated by similar post-translational mechanisms, revealing that this mechanism of determining levels of cycliln/CDK complexes is widely conserved.

In resting T cells, the “biosynthesis” of CycT1 is also regulated by miRNA-dependent inhibition of CycT1 translation in a PKC-independent manner (52,53). At this point, it is unclear whether these two apparently independent mechanisms crosstalk with each other. However, that proteasomal inhibitors increase levels of CycT1 in resting T cells to those equivalent in activated T cells suggests that protein degradation plays an essential role in maintaining extreme low levels of CycT1 in these cells. Nevertheless, disappearance of CycT1 proteins in non-replicating cells has significant clinical relevances. First this mechanism facilitates the maintenance of viral latency, including HIV, human T cell leukemia virus, cytomegalo virus, Kaposi’s sarcoma-asscociated herpes virus (KSHV) etc, which require P-TEFb for their replication (54). Moreover, our previous study shows that CycT1 is also similarly diminished in unresponsive T cells such as anergic and exhausted T cells via chronic depletion of PKC, causing P-TEFb disassembly and CycT1 degradation (27). P-TEFb is the essential co-activator of most cytokine and lymphokine genes (55). In addition, transcription factors, such as Tat, NF-kB, steroid hormone receptors, cMyc, CIITA, etc. all require P-TEFb for their effects (1,19). Therefore, maintaining high levels of CycT1, either by blocking its degradation and/or by expressing degradation-resistant CycT1 proteins in these cells could become promising approaches to improve immune function and maintain better overall cellular function.

Finally, our results provide important clues to the regulation of P-TEFb. In quiescent, terminally differentiated or anergic/exhausted cells, overall transcription levels are kept at a low level. In these cells, RNAP II is already engaged at many genes but paused at 5’ proximal regions due to the lack of P-TEFb. In actively replicating cells, P-TEFb is available and needed to transcribe genes for cell growth and proliferation. Here, equilibrium of active P-TEFb is regulated by the 7SK snRNP complex, that is essential for cellular and organismal homeostasis. Genes induced by external stimuli depend on the release of P-TEFb from the 7SK snRNP. Dysregulation of this complicated regulation of P-TEFb can result in over-expression of unwanted genes or silencing of necessary genes, as is observed commonly in many diseases and conditions. Thus, manipulating or adjusting levels of P-TEFb could become beneficial to the organism and lead to the development of new therapeutic approaches in health and diseases.

## ACKNOWLEDGEMENTS

We thank Zeping Luo and Dan Irwin (lab members) for excellent technical assistance. We thank Trang TT Nguyen in Arthur Weiss’s lab provide the cell samples. We thank Dr. Ze’ev A Ronai at Sanford Burnham Prebys Medical Discovery Institute for valuable suggestions and helpful discussions.

